# Molecular phylogeny of *Odontosyllis* (Annelida, Syllidae): A recent and rapid radiation of marine bioluminescent worms

**DOI:** 10.1101/241570

**Authors:** Aida Verdes, Patricia Alvarez-Campos, Arne Nygren, Guillermo San Martin, Greg Rouse, Dimitri D. Deheyn, David F. Gruber, Mande Holford

## Abstract

Marine worms of the genus *Odontosyllis* (Syllidae, Annelida) are well known for their spectacular bioluminescent courtship rituals. During the reproductive period, the benthic marine worms leave the ocean floor and swim to the surface to spawn, using bioluminescent light for mate attraction. The behavioral aspects of the courtship ritual have been extensively investigated, but little is known about the origin and evolution of light production in *Odontosyllis*, which might in fact be a key factor shaping the natural history of the group, as bioluminescent courtship might promote speciation. To investigate the speciation patterns and evolutionary history of *Odontosyllis* and to trace the origin of bioluminescence within the group, we inferred phylogenies using both gene concatenation and multispecies coalescent species-tree approaches with a multilocus molecular dataset (18S *rRNA*, 16S *rRNA* and COI). We also used this dataset to estimate divergence times and diversification rates in a relaxed molecular clock Bayesian framework. Our results suggest that *Odontosyllis* has undergone a recent rapid radiation, possibly triggered by the origin of bioluminescent courtship, which might have increased speciation rates and lineage divergence through sexual selection. Additionally, our analyses reveal that the genus *Odontosyllis* as currently delineated is a paraphyletic group that needs to be reorganized to reflect evolutionary relationships.

## 1. Introduction

The genus *Odontosyllis* is a lineage of marine annelids that has sparked the curiosity of many scientists during the last century because of its spectacular bioluminescent courtship displays (Figure 1) (Crawshay, 1935; Deheyn and Latz, 2009; Erdman, 1965; Fischer and Fischer, 1995; Gaston and Hall, 2000; Huntsman, 1948; Markert et al., 1961; Potts, 1913; Shimomura et al., 1963; Tsuji and Hill, 1983). *Odontosyllis* belongs to the family Syllidae, a diverse group of marine annelids present in nearly all marine benthic habitats and characterized by the proventricle, a specialization of the digestive tube (Aguado et al., 2012; Rouse and Pleijel, 2001). The taxonomy and systematics of syllids is considerably complicated for several reasons. Arguably the most significant one, is that numerous key morphological characters traditionally used to identify species are homoplastic, and in many occasions they can been misinterpreted leading to erroneous species identifications (Álvarez-Campos et al., 2017a; Álvarez-Campos and Verdes, 2017). Additionally, although so far only two cases have been confirmed, pseudo-cryptic speciation might be common among syllids (Álvarez-Campos et al., 2017a, 2017b). As a result, the taxonomic status of many lineages is not clear and the current classification of numerous genera needs to be revised and ideally based on studies that combine thorough morphological and molecular analyses (Álvarez-Campos et al., 2017b). The genus *Odontosyllis* is one of the various syllid lineages in need of a detailed systematic revision.

**Figure 1.**
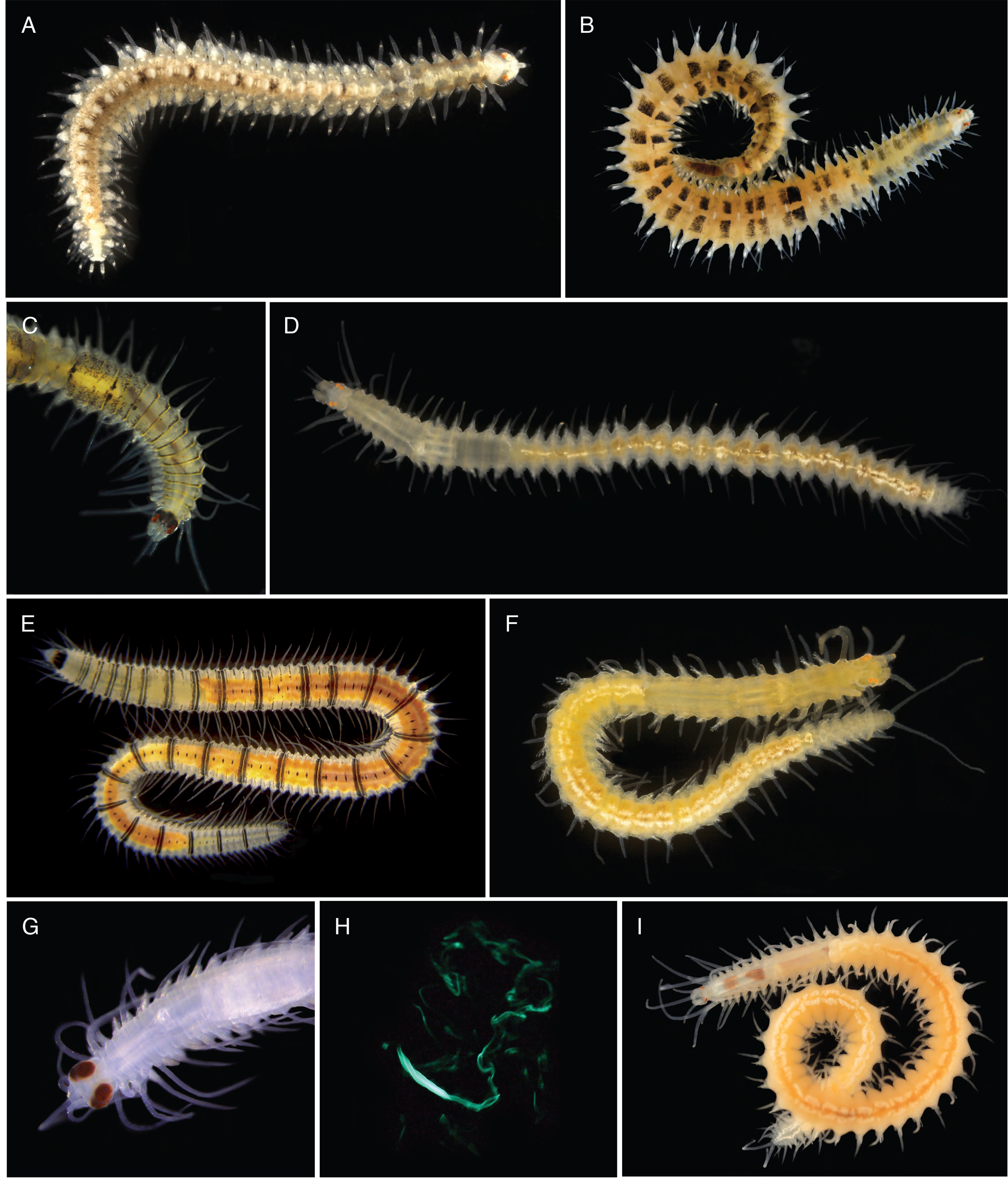
Images of *Odontosyllis* and *Eusyllis* species. (A) *Odontosyllis gibba*, type species of the genus; (B) *Odontosyllis fasciata*; (C) *Odontosyllis phosphorea*, anterior end; (D) *Odontosyllis* sp.; (E) *Odontosyllis suteri*; (F) *Odontosyllis* sp.; (G) *Odontosyllis enopla* male, anterior end; (H) Bioluminescent display of female of *Odontosyllis enopla*; (I) *Eusyllis blomstrandi*, closely related bioluminescent species. Images courtesy of Arne Nygren (A, D–F), Fredrick Pleijel (B, I) and John Sparks (H).

*Odontosyllis* species are characterized by a combination of morphological characters including the presence of a well-developed occipital flap covering the posterior part of the prostomium (Figure 1) and a short pharynx with a few teeth directed backwards and flanked by two lateral plates (Fukuda et al., 2013; San Martin and Hutchings, 2006; Verdes et al., 2011). This unique pharyngeal armature referred to as a denticled arc, is considered the main diagnostic feature of the genus (San Martin and Hutchings, 2006). It might seem straightforward to identify *Odontosyllis* species based on this distinctive morphological characteristic, but many of the 55 species that currently comprise the genus (WoRMS Editorial Board, 2017) correspond to single reports whose descriptions lack crucial information regarding the structure of the pharyngeal armature, hindering any revisionary efforts (Fukuda et al., 2013; Verdes et al., 2011). Additionally, identification to the species level is significantly difficult since many species have been described using different combinations of characters, and in several cases the descriptions are based on slight morphological variations (Fukuda et al., 2013).

Several *Odontosyllis* species show an impressive bioluminescent display during reproduction, that has captivated humans since Christopher Columbus approached the Bahamas in 1492 (Crawshay, 1935). Bioluminescence has evolved independently in several annelid lineages (Haddock et al., 2010; Verdes and Gruber, 2017), including remarkable species like the deep sea *Swima bombiviridis* that releases luminescent “bombs” (Osborn et al., 2009) or the pelagic tomopterids that emit yellow light, an extremely rare color among marine luminescent taxa (Widder, 2010). There are many other examples among the 98 luminous annelids reported so far, which occupy a variety of habitats from terrestrial to marine ecosystems, and display a wide range of bioluminescent colors associated with different functions, such as defense or intraspecific communication (Verdes and Gruber, 2017). In *Odontosyllis*, bioluminescent light is used for mate attraction in adults, acting as a swarming cue during reproduction. In the summer months, a few days after a full moon and shortly after sunset, the benthic marine worms leave the ocean floor and swim to the surface to reproduce. In most cases, females appear first and swim in circles while releasing gametes in a cloud of bright green luminous mucus that attracts the males, which dart towards the females to spawn (Galloway and Welch, 1911; Markert et al., 1961; Verdes and Gruber, 2017). This courtship ritual has been thoroughly described from a behavioral point of view (Fischer and Fischer, 1995; Gaston and Hall, 2000; Huntsman, 1948; Markert et al., 1961; Tsuji and Hill, 1983), but little is known about the basic chemistry of the bioluminescence system or the evolution of light production in *Odontosyllis*. Additionally, a recent study by Ellis and Oakley (2016) showed that lineages with bioluminescent courtship displays (including syllids) are more speciose and have significantly higher rates of diversification than their non-luminous sister groups. Because bioluminescence courtship is almost certainly a sexually selected trait, this study supports the longstanding theory that sexual selection promotes speciation at a macroevolutionary scale (Ellis and Oakley, 2016; Kraaijeveld et al., 2011).

Thus, *Odontosyllis* is a very promising group to investigate a variety of questions ranging from the systematics of conflictive taxa, to the implications of key innovations such as bioluminescence in the speciation process. Nevertheless, the inference of species boundaries and phylogenetic relationships is fundamental for any systematic, ecological or evolutionary study and consequently, a robust phylogenetic framework of the genus *Odontosyllis* is a necessary backbone to explore any of the aforementioned questions. To clarify the evolutionary history of *Odontosyllis* and the closely related *Eusyllis*, we inferred phylogenies using DNA sequence data from nuclear marker 18S rRNA and mitochondrial markers 16S rRNA and cytochrome oxidase subunit I from 63 *Odontosyllis* specimens, 13 *Eusyllis* and 35 outgroups. We followed both a gene concatenation approach using maximum likelihood (ML) and Bayesian inference (BI) analyses and a multispecies coalescent species-tree approach. We also used the concatenated dataset in a relaxed molecular clock Bayesian analysis to simultaneously estimate topology, divergence times and diversification rates. We discuss the taxonomical implications of our results and explore possible scenarios for the origin and evolution of light production and the role of bioluminescence courtship as a driver of speciation in *Odontosyllis*.

## 2. Material and methods

### 2.1. Sample collection

Samples were collected in different biological surveys between 2012 and 2016 from a variety of substrates in intertidal and subtidal zones by hand, snorkeling or SCUBA diving. Information regarding collection dates, locality and substrates is listed in Supplementary Table S1. Specimens were sorted in the field using light microscopy and fixed in 96% ethanol or RNAlater for morphological and molecular analyses. Prior to fixation, selected specimens were anaesthetized with 7% magnesium chloride buffered in seawater and photographed under a microscope. Further examination and identification was completed using a Nikon Optiphot light microscope with a differential interference contrast system (Nomarsky) at Universidad Autónoma de Madrid (UAM).

### 2.2. DNA extraction, amplification and sequencing

Genomic DNA was extracted using the DNeasy Blood & Tissue Kit (Qiagen), following manufacturer’s protocols. Fragments of the nuclear gene *18S rRNA* (1720 bp), and the mitochondrial *16S rRNA* (609 bp) and cytochrome c oxidase subunit I (COI, 659 bp) were amplified by polymerase chain reaction (PCR). Three overlapping pairs of primers were used to amplify *18S rRNA*: 18S1F-18S4R, 18S3F-18Sbi, and 18Sa2.0-18S9R (Giribet et al., 1996). Primers 16SarL and 16SbrH (Palumbi, 1996) were used to amplify 16S rRNA, and the modified primers with inosine jgLCO1490 and jgHCO2198 (Geller et al., 2013) were employed to amplify COI in all specimens. The PCR reactions consisted of 1 μL of DNA template in 25 μL reaction volumes containing 18 μL H2O; 5 μL of 59 USB buffer, 0.25 μL of each of 10 μM primers, 0.5 μL of 10 mM dNTPs, and 0.13 μL of 1.25 U/μL GOTaq DNA Polymerase (Promega). The temperature profile for the 18S and 28S rRNA nuclear markers was as follows: 95 °C/120 s; (95 °C/30 s; 47 °C/30 s; 72 °C/180 s) 9 35 cycles; 72 °C/300 s; for 16S rRNA: 95 °C/5 min; (95 °C/30 s; 45 °C/30 s; 72 °C/60 s) 9 35 cycles; 72 °C/10 min; and for COI: 95 °C/15 min; (94 °C/30 s; 45 °C/70 s; 72 °C/90 s) 9 40 cycles; 72 °C/10 min. 1.5 µL of the PCR product was used for sequencing using the forward primer of the primers described above at the Servicio de Secuenciación Sanger, Unidad de Genómica (Universidad Complutense de Madrid) and at Genewiz Inc. (South Plainfield, NJ).

Sequences were edited in Geneious v8.1.9 (Kearse et al., 2012), to remove primers from the three *18S rRNA* fragments and merge the overlapping fragments into a consensus sequence. Multiple sequence alignments for the three different genes were built in the online server of MAFFT v7 under default parameters (Katoh and Standley, 2013). To remove poorly aligned positions and highly variable regions in the *18S rRNA* gene, sequences were run in Gblocks v0.91b (Castresana, 2000) with all options for a less stringent selection selected.

### 2.3. Phylogenetic analyses

#### 2.3.1. Gene concatenation approach: ML (RAxML) and BI (MrBayes and BEAST)

In order to assess the monophyly of *Odontosyllis* and the closely related *Eusyllis*, we analyzed the three molecular markers (18S, 16S and COI) in a total of 111 specimens (63 *Odontosyllis*, 13 *Eusyllis* and 35 outgroups) following a gene concatenation approach using both maximum likelihood (ML) and Bayesian inference (BI). All mitochondrial and nuclear data sets were concatenated and the best-fitting model of sequence evolution was selected using the Akaike information criterion (AIC) in jModeltest 2 (Darriba et al., 2012). The best model for the concatenated data set was a general time-reversible (GTR) with gamma-distributed rates across sites and a proportion of invariable sites (GTR + G + I). Partitions for each of the markers were used in all subsequent phylogenetic analyses: for ribosomal markers, non-codon-specific models were used; and for COI, we used codon-specific models. ML analyses were run in RAxML v1.31 (Stamatakis, 2006) using the GTR+G+I evolutionary model. Bootstrap support values were estimated using 1000 replicates and 10 starting trees (Stamatakis et al., 2008). BI analyses were run with MrBayes v3.2.1 (Ronquist et al., 2012) in the CIPRES Science Gateway v3.1 (Miller et al., 2010). Analyses were run with using the GTR+G+I evolutionary model, with four Markov chains that were started from a random tree, running simultaneously for 100 million generations, with trees sampled every 10,000 generations (samplefreq = 10000); the initial 25 % of trees were discarded as burnin (burninfrac=0.25) after assessing for convergence with Tracer v1.6.0 (Rambaut and Drummond, 2013).

#### 2.3.2. Multispecies coalescent species-tree approach

Preliminary analyses suggested topological discordance between mitochondrial and nuclear phylogenies, which might reflect the presence of incomplete lineage sorting (ILS). To account for ILS we inferred the species tree under a Bayesian multispecies coalescent framework using *BEAST (StarBeast) (Heled and Drummond, 2010) as implemented in BEAST2 v2.4.6 (Bouckaert et al., 2014). This method coestimates multiple gene trees embedded in a shared species tree, explicitly incorporating the coalescent process and the expected discordance among genes resulting from ILS (Heled and Drummond, 2010; Townsend et al., 2011). Separate and unlinked substitution models were defined for each gene, corresponding to the best-fitting model of sequence evolution selected by jModeltest 2 (Darriba et al., 2012). Linked tree and clock models were specified for the mitochondrial genes whereas separate unlinked tree and clock models were defined for the nuclear marker. We used an uncorrelated lognormal relaxed molecular clock and a strict molecular clock for the mitochondrial and nuclear genes respectively, with relative rates estimated and a Yule model as the species tree prior. Divergence time estimates were constrained by placing age calibrations priors as lognormal distributions on two nodes, based on fossil data (Phyllodocida = 485 ± 1.9 Ma and Goniadidae = 323 Ma) (Parry et al., 2014). The *BEAST analysis ran for 500,000,000 generations sampling every 50,000 and the first 10% of samples were discarded as burnin. We used Tracer (Rambaut and Drummond, 2013) to assess convergence of parameter estimates and to verify that effective sample size (ESS) values were above 200 for all parameters after burnin was discarded. The resulting tree topologies, branch lengths and divergence times were summarized on a maximum clade credibility tree using TreeAnnotator v2.4.7 (Bouckaert et al., 2014).

### 2.4. Divergence dating and diversification rate estimation

To determine divergence times in syllid lineages we used the three loci concatenated dataset and run a Bayesian uncorrelated relaxed clock analysis with two fossil calibrations (as defined above) in BEAST 2 v2.4.6 (Bouckaert et al., 2014). Evolutionary rates along branches followed an uncorrelated lognormal relaxed molecular clock for the mitochondrial markers and a strict molecular clock for the nuclear gene, with relative rates estimated and a Yule model as the species tree prior. The BEAST analysis ran for 300,000,000 generations sampling every 30,000 and the first 10% of samples were discarded as burnin. Tracer (Rambaut and Drummond, 2013) was used to assess convergence and confirm ESS values were at least 200 for all estimated parameters after burnin was removed. The resulting tree topologies, branch lengths and divergence times were summarized on a maximum clade credibility tree using TreeAnnotator v2.4.7 (Bouckaert et al., 2014).

### 2.5. Ancestral State Character Reconstruction

To investigate the evolution of bioluminescence in the group we performed ancestral state reconstructions (ASR) on both time-calibrated ultrametric phylogenies generated, the BEAST dated tree and the species tree topologies. Bioluminescence was coded as a binary character (0: absence, 1: presence) for the species for which information was available and ASR was performed in Mesquite v3.31 (Maddison and Maddison, 2016) using equally weighted, unordered states. ML optimization was implemented under the Mk1 model, to identify the state at each node that maximizes the probability of the states observed in the terminal taxa (Pagel, 1999). Outgroup representatives and species duplicates were excluded from the analyses.

## 3. Results

### 3.1. Phylogenetic analyses

#### 3.1.1. Gene concatenation approach

Final multiple sequence alignments included partial sequences of nuclear marker *18S rRNA* (1720 bp after Gblocks) from 101 specimens, and mitochondrial genes *16S rRNA* from 76 specimens (609 bp) and COI from 86 specimens (659 bp). The best fitting model of sequence evolution for both nuclear and mitochondrial markers under AIC was the GTR + G + I. Both ML (Figure 2) and BI (Supplementary Figure S2) analyses of the three concatenated genes recovered a paraphyletic *Odontosyllis*, as *Eusyllis kupfferi* and *E. blomstrandi* are found nested within *Odontosyllis* (Figure 2). Since the two topologies obtained from the ML and BI analyses are almost identical, we present the ML tree with both bootstrap (BS) and posterior probabilities (PP) support values mapped on each node (Figure 2).

**Figure 2.**
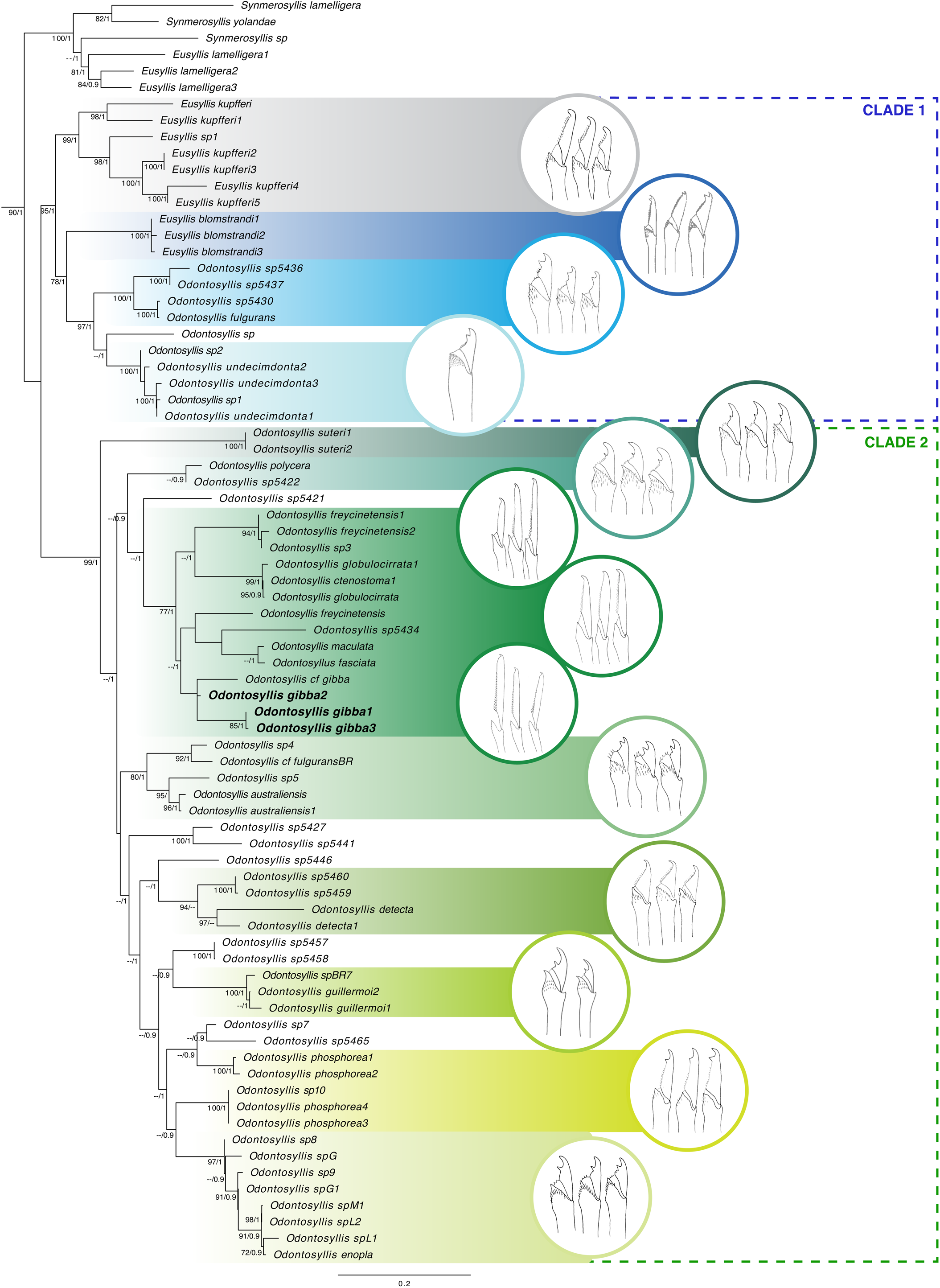
Phylogenetic relationships of *Odontosyllis* and the closely related *Eusyllis* inferred from the ML analysis of the concatenated dataset (*18S rRNA*, *16S rRNA* and *COI*). The characteristic chaetae of supported clades are shown to the right. The type species *O. gibba* is marked in bold. Numbers below branches indicate bootstrap support values (BS > 75%) and posterior probabilities (PP > 0.90). Eusyllinae outgroups removed for clarity.

Our analyses recovered a paraphyletic *Odontosyllis*, with two main well-supported clades labeled Clade 1 and Clade 2 in Figure 2. Clade 1 includes *O. fulgurans*, *O. undecimdonta*, three unidentified *Odontosyllis* species and two *Eusyllis* species: *Eusyllis kupfferi* and *E. blomstrandi*. Clade 2 groups all remaining *Odontosyllis* and includes the type species *O. gibba* (Figure 2). Within Clade 1, both analyses show strong support for a lineage that includes *O. fulgurans*, *O. undecimdonta* and three unidentified *Odontosyllis*. Within Clade 2, both analyses recovered a well-supported clade including the type species *O. gibba* and other species characterized by unidentate chaetae with long blades, including *O. freycinetensis*, *O. globulocirrata*, *O. maculata* (Figure 2). The clade comprising *O. australiensis*, *O. cf. fulgurans* and two unidentified species from Brazil and Indonesia (*Odontosyllis* sp4 and *Odontosyllis* sp5 respectively) was also well supported in both analyses along with a clade including *O. enopla* and seven undescribed Caribbean *Odontosyllis* (Figure 2).

There were slight differences in the topologies recovered in the ML and BI analysis with regards to the relationships among species of Clade 2. In particular, the clade comprising *O. australiensis*, *O. cf. fulgurans* and two unidentified species appears more closely related with the group of species with short bidentate chaetae in the ML tree (Figure 2) while in the BI topology this clade is more closely related with the group of species with unidentate chaetae and long blades (Supplementary Figure S2). Additionally, in the ML tree the clade including GenBank sequences from *O. phosphorea* and two unidentified *Odontosyllis* appears closely related with a clade including other *O. phosphorea* specimens, *O. enopla* and undescribed Caribbean *Odontosyllis*, whereas in the BI topology it appears more closely related with a group including *O. guillermoi*, and *Odontosyllis* sp4. Nevertheless, the conflicting nodes are weakly supported in both analyses and the overall results are almost identical.

#### 3.1.2. Multispecies coalescent species-tree approach

The topology of the Bayesian species-tree (Figure 3) is almost identical to that from the ML and BI analyses of the concatenated data (Figure 2 and Supplementary Figure S2). In contrast to the concatenation approach however, ILS is specifically modeled in the species-tree estimation. The topologies differ only in the position of a handful of species, including *Eusyllis kupfferi* and *Eusyllis* sp1 which in the ML and BI trees appear nested in Clade 1 and are sister to the lineage that includes the species *O. fulgurans* (Figure 2), whereas in the species-tree they are found within Clade 2 (Figure 3). The position of *O. suteri* is also different, being placed as the sister species to all other Clade 2 taxa in the ML and BI trees (Figure 2), whereas in the species-tree is found as the sister lineage of the group of *Odontosyllis* with short bidentate blades (Figure 3). The topologies also differ in the placement of *Odontosyllis* sp7, which in the species-tree is more closely related to *Odontosyllis cf. fulgurans*, and *Odontosyllis* sp4 (Figure 3), while in the ML and BI trees is nested with *O. enopla* and other undescribed Caribbean *Odontosyllis* (Figure 2).

**Figure 3.**
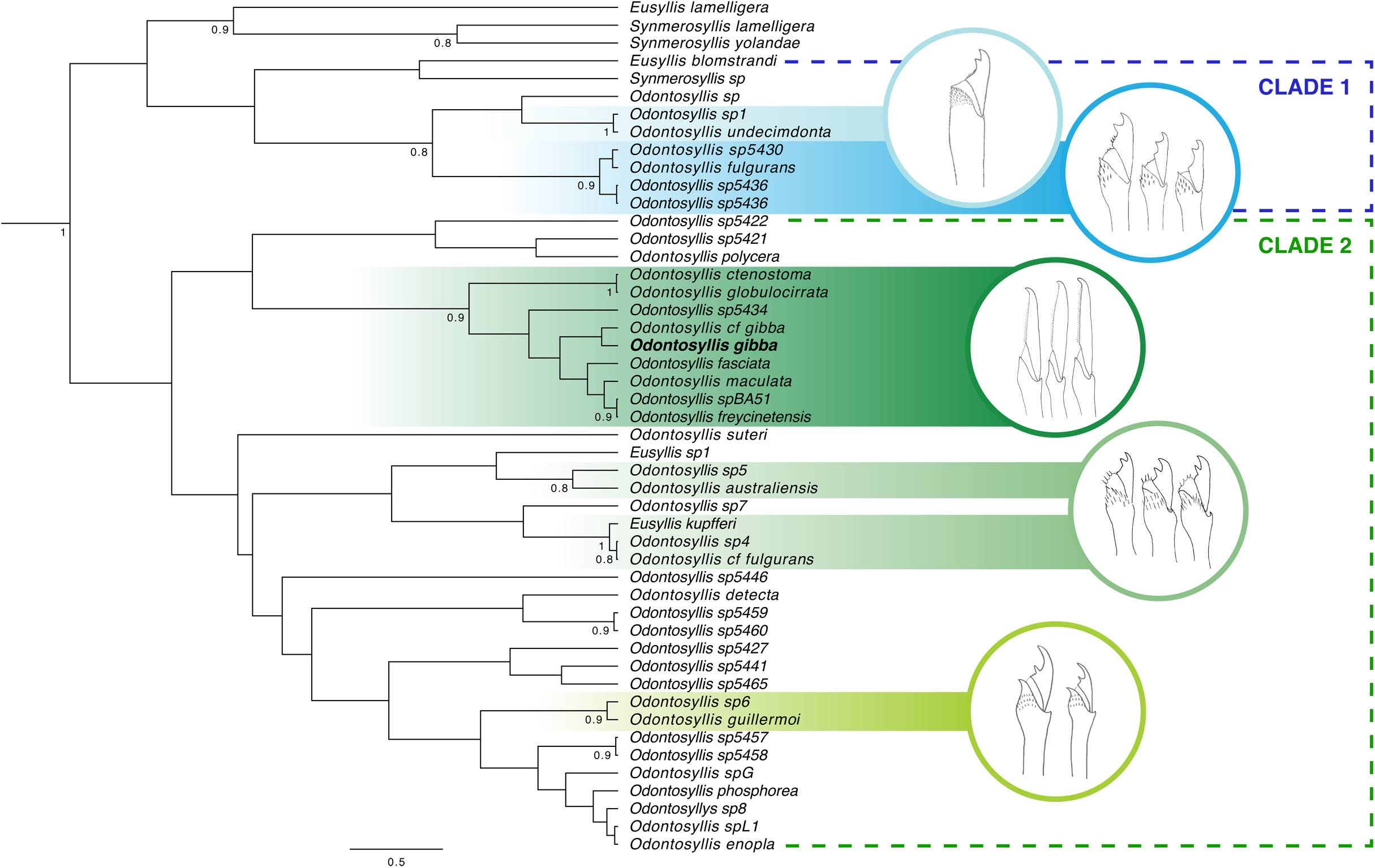
Bayesian multispecies coalescent species-tree of *Odontosyllis* and the closely related *Eusyllis*. The characteristic chaetae of supported clades are shown to the right. The type species *O. gibba* is marked in bold. Numbers below branches indicate posterior probabilities (PP > 0.80). Eusyllinae outgroups removed for clarity.

Despite some topological differences with the ML and BI trees, the two main clades (Clade 1 and Clade 2) are also recovered in the species-tree, although these groups are not well supported (Figure 3). The clade including the species *O. fulgurans*, *O. undecimdonta* and three unidentified *Odontosyllis* is also recovered with weak support, whereas the clade comprising the type species *O. gibba* and the rest of *Odontosyllis* with unidentate chaetae and long blades, including *O. freycinetensis*, *O. globulocirrata* and *O. maculata* is also recovered with strong support (Figure 3).

### 3.2. Divergence dating and diversification rate estimation

The BEAST2 time-calibrated tree topology is almost identical to that recovered in the ML and BI analyses, although the support is increased for almost all nodes (Figure 4 and Supplementary Figure S3). In this Bayesian analysis of the concatenated dataset, we estimated divergence times for the four Syllidae subfamilies along with several Eusyllinae lineages. Our results suggest that the family Syllidae is an old lineage of annelids that appears to have diverged in the Ordovician-Silurian boundary during the Paleozoic (Figure 4). Out of the four syllid subfamilies, Syllinae appears to be the oldest lineage, with an estimated divergence during the Silurian (425.9 Ma), followed by Autolytinae which is estimated to have diverged during the Triassic (235.9 Ma) and Eusyllinae, during the Jurassic (166.6 Ma). The subfamily Exogoninae seems to be the youngest syllid lineage, having diverged during the Cretaceous (83.44 Ma) (Figure 4).

**Figure 4.**
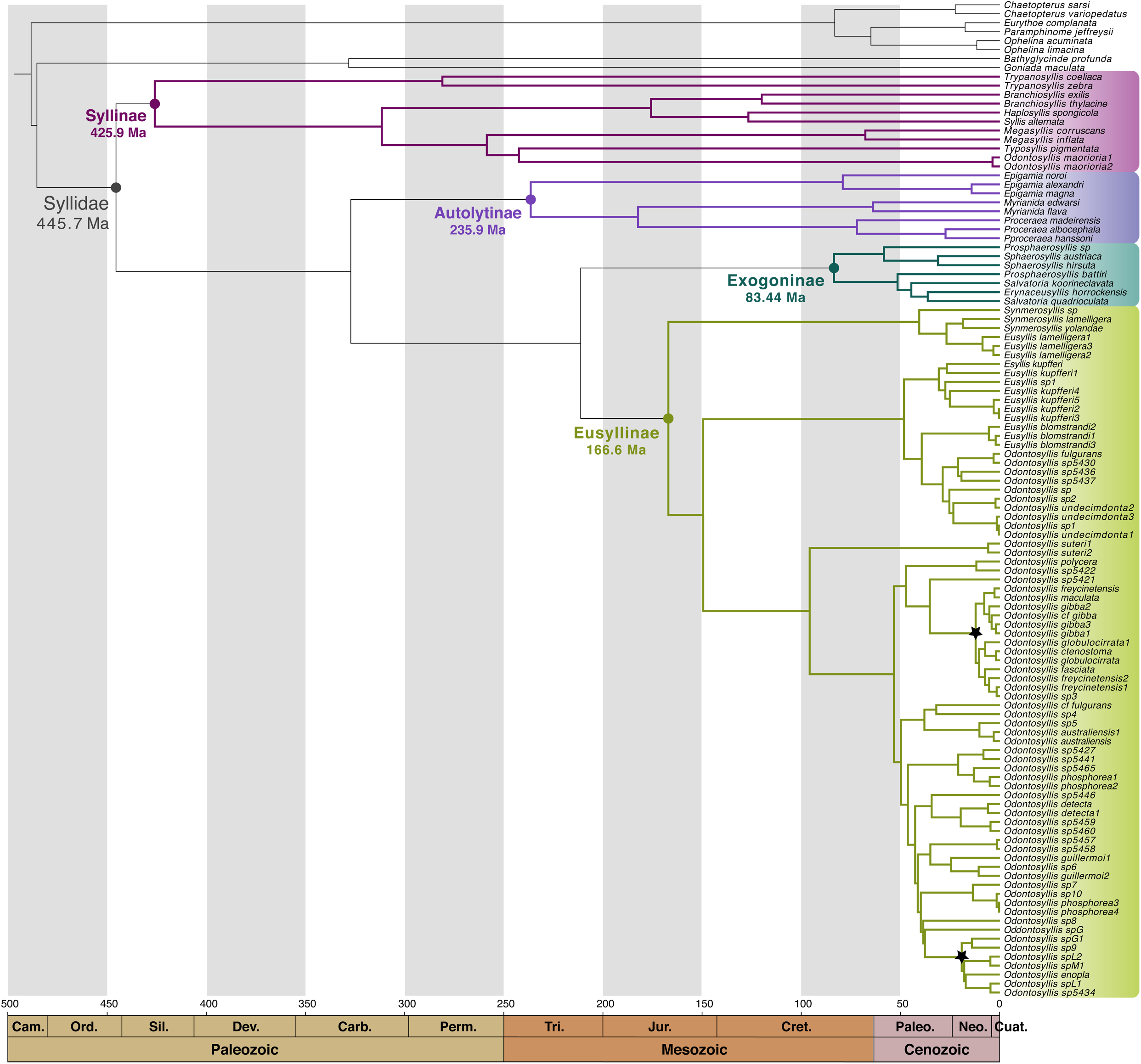
Time-calibrated phylogeny of major lineages of Syllidae inferred through Bayesian analyses of the concatenated dataset. Estimated divergence dates for the four subfamilies are specified in millions of years (Ma) and the corresponding geological periods are indicated at the bottom. Asterisks denote two recent and rapid speciation events in *Odontosyllis*.

Interestingly, the speciation process in *Odontosyllis* and other Eusyllinae genera appears to be very recent in comparison with genera from other syllid subfamilies, including Exogoninae. For instance, while many *Odontosyllis* species appear to have diverged in the Neogene, during the Cenozoic (asterisks in Figure 4), *Trypanosyllis* (Syllinae) species diverged in the Permian, whereas *Branchiosyllis* and *Megasyllis* (Syllinae) diverged in the Cretaceous (Figure 4). Species of the Autolytinae genera *Myrianida*, *Epigamia* and *Proceraea* appear to also have diverged in the Cretaceous, during the Mesozoic (Figure 4). Although the subfamily Exogoninae is estimated to have diverged more recently than Eusyllinae, species diversification seems to have started earlier than in Eusyllinae genera, specifically during the Paleogene (Figure 4). However, this last result is unclear due to apparent non-monophyly of the Exogoninae genera included in the analysis.

In addition, the time-calibrated phylogeny allowed the implementation of an uncorrelated, lognormal relaxed clock model to estimate the branch-specific substitution rates among Eusyllinae lineages (Figure 5). The branches of the time-calibrated phylogeny are colored according to the estimated diversification rates in number of substitutions per site per year in Figure 5. Interestingly, the majority of the highest diversification rates are associated with lineages that include bioluminescent species. For instance, the highest rate was estimated to be 1.45E-7 substitutions/site/year and is associated with a clade that includes *O. enopla*, *O. phosphorea* and several undescribed Caribbean *Odontosyllis* that use bioluminescence for courtship (Figure 5).

**Figure 5.**
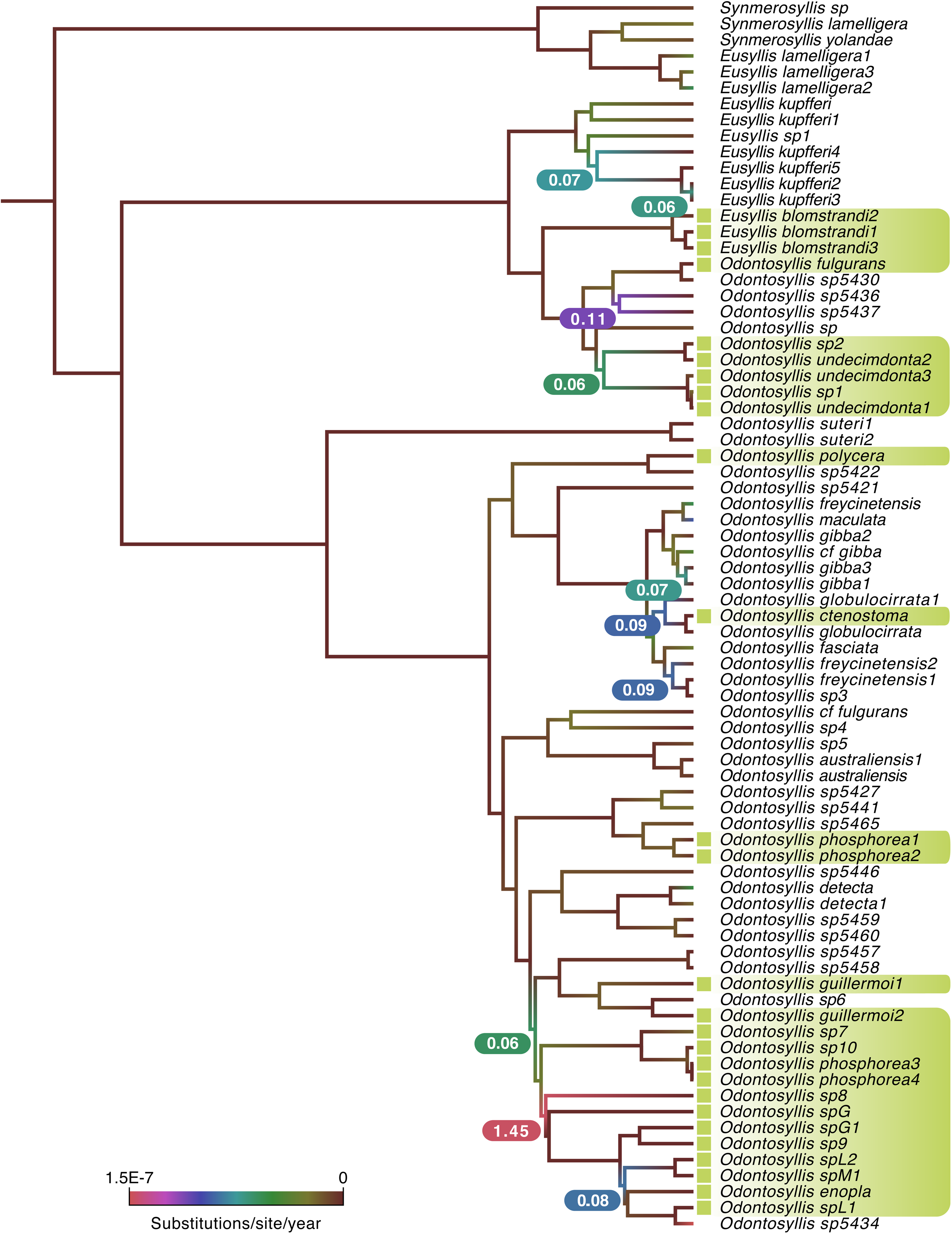
Diversification rates in Eusyllinae lineages. Time-calibrated phylogeny with branches colored according to substitution rate (substitution per site per year). Highest diversification rates are shown next to the corresponding node (in substitutions per site per 10 million years). Bioluminescent species are highlighted in green and Eusyllinae outgroups are removed for clarity.

### 3.3. Ancestral state character reconstruction

The results from both ASR analyses using the *BEAST multispecies coalescent species-tree and the BEAST2 time-calibrated tree, support that bioluminescence was the ancestral state, with evidence suggesting it was present in the most recent common ancestor of extant Eusyllinae lineages (Figure 6). Both ML reconstructions suggest with high probability (0.73 in BEAST2 tree and 0.71 in species-tree) that bioluminescence evolved once within Eusyllinae, with secondary losses in several lineages. With regard to *Odontosyllis* both ASR analyses also show strong support for a single origin of bioluminescence, with a high probability that the ability to produce light was the ancestral state for the group (0.71 in the species-tree and 0.83 in the BEAST2 tree) (Figure 6).

**Figure 6.**
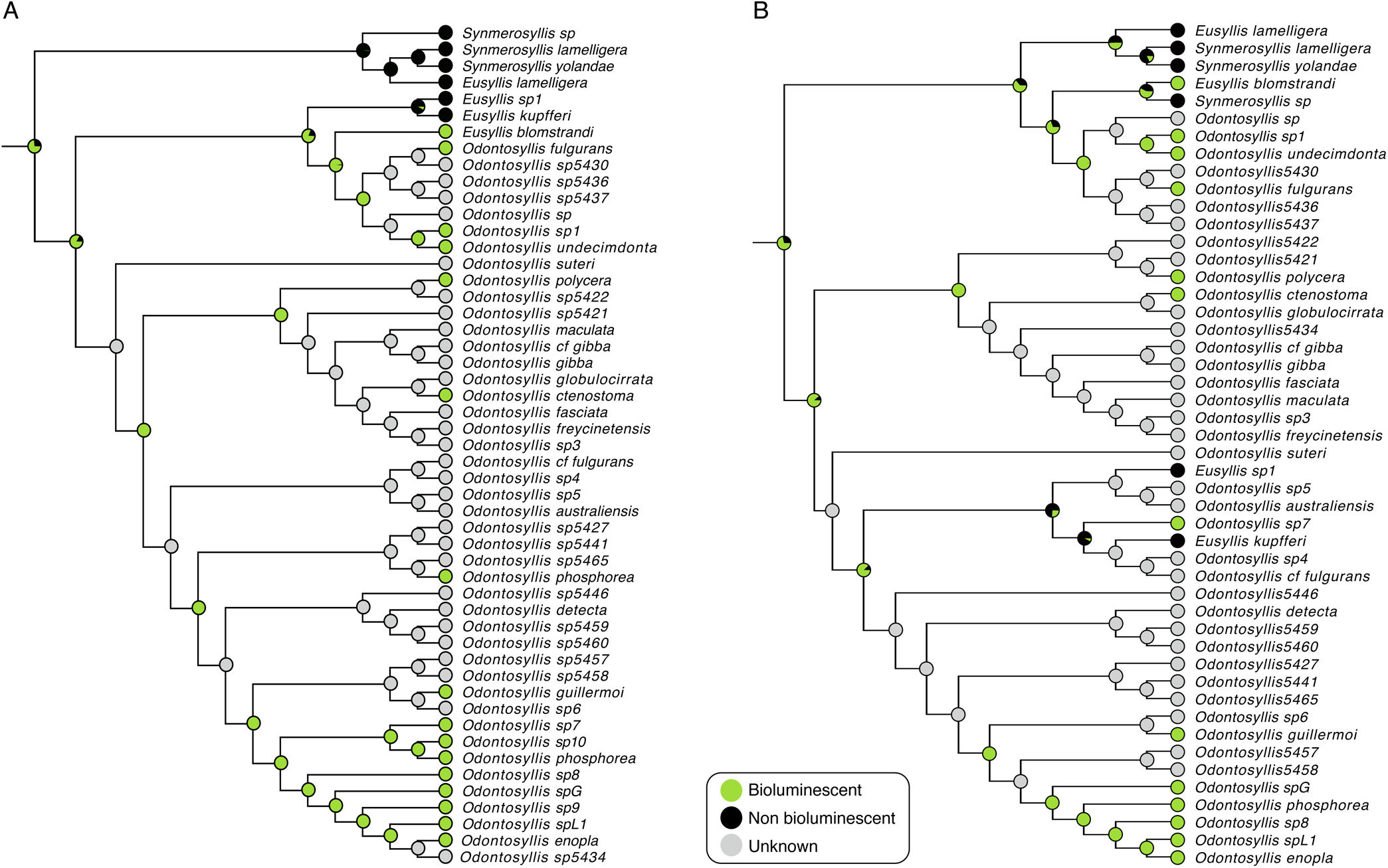
Maximum likelihood ancestral state reconstruction of bioluminescence in Eusyllinae lineages. (A) Reconstruction based on the BEAST2 time-calibrated phylogeny; (B) Reconstruction using the multispecies coalescent species-tree. Outgroups and species duplicates are excluded from the analyses.

## 4. Discussion

### 4.1. Phylogenetic analyses and taxonomical implications

Preliminary analyses of the molecular data suggested topological discordance between mitochondrial and nuclear phylogenies, which can indicate the presence of incomplete lineage sorting (ILS), a process by which ancestral polymorphisms are retained through speciation events, and might compromise concatenated phylogenetic analyses (Philippe et al., 2011; Townsend et al., 2011) (Townsend et al., 2010). Thus, in addition to gene concatenation, we used Bayesian multispecies coalescent methods to coestimate multiple gene trees embedded in a shared species tree, explicitly incorporating the expected discordance among gene trees resulting from ILS (Heled and Drummond, 2010; Townsend et al., 2011).

All the phylogenetic analyses completed in this study recovered a paraphyletic *Odontosyllis*, with some species of the closely related genus *Eusyllis* nested within (Figures 2 and 3). Despite some conflictive nodes, all methods recovered two main clades, Clade 1 and Clade 2, with strong support in the gene concatenation analyses (Figure 2) but not well supported in the species-tree (Figure 3). Within Clade 2, all analyses recovered a well-supported lineage that includes *O. freycinetensis*, *O. cf. gibba*, *O. globulocirrata*, *O. maculata* and the type species *O. gibba* and thus, this lineage would represent the taxonomically valid *Odontosyllis* (Figures 2 and 3). Initial morphological examination showed that this Clade 2 lineage is comprised by species with unidentate chaetae and long blades, not found in other lineages.

Although the resulting topologies are overall similar, they differ in the placement of some species and the support values are increased in the trees derived from the gene concatenation approach. This topological incongruences between gene concatenation and species-tree methods and the low support values in the species tree are indicative of conflicting gene topologies, likely due to ILS. Gene concatenation analyses showed stronger support than species-tree methods, however, ILS is not considered in analyses of concatenated data and can result in inconsistent phylogenetic estimates and high support for an incorrect topology (Kubatko and Degnan, 2007; Kutschera et al., 2014). Species-tree methods on the other hand, explicitly incorporate the expected discordance among genes and have been shown to provide more accurate estimates of the species tree than concatenation analyses (Heled and Drummond, 2010; Lambert et al., 2015; Maddison and Knowles, 2006). Unfortunately, our species-tree shows weak support in most nodes and consequently, given the inconsistencies of the phylogenetic analyses and the lack of distinctive morphological features identified in the supported lineages, we do not consider appropriate taking any taxonomic action until new material is available for molecular study and more thorough morphological analyses can be completed.

### 4.2. Divergence dating analyses

Our time-calibrated phylogeny suggests that Syllidae is an ancient group of annelids that must have diverged in the Ordovician-Silurian boundary during the Paleozoic (Figure 4). This estimation would place the divergence of syllids approximately 50 million years before the oldest unequivocal phyllodocidan fossils known, which correspond to the Middle Devonian (Farrell and Briggs, 2007) and much older than the first appearance of many phyllodocidan families in the Carboniferous (Parry et al., 2014). However, Parry et al. (2014) suggest that this collection of phyllodocidan fossils from the Carboniferous is likely due to favorable taphonomic conditions rather than a true Carboniferous radiation of phyllodocidans.

The Eusyllinae lineage is estimated to have diverged during the Jurassic and it appears to have radiated into several lineages during the Cretaceous (Figure 4), including the *Odontosyllis* lineages identified as Clade 1 and Clade 2 in the phylogenetic analyses. The speciation process in the Eusyllinae lineages seems to be more recent than in lineages from other syllid subfamilies, and *Odontosyllis* specifically, seems to have split into several lineages over a relatively short period of time (see short internal branches in Figure 4). This pattern of rapid branching, characteristic of speciation events closely spaced in time, has probably resulted in incomplete lineage sorting (Townsend et al., 2011). This is consistent with our phylogenetic results, which show a lack of phylogenetic signal, leading to short internal branches that are difficult to resolve (Philippe et al., 2011).

### 4.3. Origin and evolution of bioluminescence in *Odontosyllis*

It is well known that many species of *Odontosyllis* are luminous (Figures 1A, C, G, H), but there are also reports of bioluminescent species in other Eusyllinae genera including *Eusyllis* (Figure 1I) and *Pionosyllis* (Haddock et al., 2010; Verdes and Gruber, 2017; Zörner and Fischer, 2007). The ancestral state reconstruction analyses we propose tracing the evolution of bioluminescence in Eusyllinae suggest that the capability to produce light evolved once within the group and was present in both the most recent common ancestor of extant Eusyllinae lineages, and the most recent common ancestor of *Odontosyllis* (Figure 6).

Several *Odontosyllis* species display a bioluminescence courtship ritual, in which light is used for mate attraction and functions as a swarming cue during reproduction (Gaston and Hall, 2000; Tsuji and Hill, 1983; Verdes and Gruber, 2017). A recent study showed that linages with bioluminescent courtship displays are associated with higher species richness and faster rates of species accumulation than non-luminous relatives (Ellis and Oakley, 2016). The time-calibrated phylogeny allowed us to estimate the lineage-specific speciation rates in luminous syllids, showing that faster rates of diversification are associated with lineages that include bioluminescent species (Figure 5). For some *Odontosyllis* species such as *O. ctenostoma* or *O. polycera* there is only a handful of reports documenting the capability of the worms to glow, and whether they use bioluminescence for courtship is not clear. However, the fastest speciation rate estimated in our analyses (1.45E-7 substitutions/site/year) corresponds to a lineage that includes *O. phosphorea* and *O. enopla*, two species with well-known luminous courtship rituals, in addition to other Caribbean undescribed species that also use light for courtship (personal observation) (Figure 5). Our results suggest that the origin of bioluminescent courtship might be associated with an increase in speciation rates, potentially leading to higher species richness in luminous courting syllids. Our findings are in line with Ellis and Oakley (2016) and provide further support to the theory that sexual selection increases speciation rates at a macroevolutionary scale (Ellis and Oakley, 2016; Kraaijeveld et al., 2011).

## 5. Conclusion

In conclusion, *Odontosyllis* seems to have undergone a recent rapid radiation, possibly triggered by the origin of bioluminescent courtship, which would increase speciation rates and lineage divergence through sexual selection. This rapid radiation has led to ILS, causing topological incongruences between gene trees, and between trees inferred from gene concatenation and coalescent-based methods, as well as low support values in the species-tree.

Despite these inconsistencies, all analyses clearly show that *Odontosyllis* as currently delineated is a paraphyletic group and needs to be reorganized to reflect evolutionary relationships. However, our results also indicate that the genetic markers employed have low phylogenetic signal, leading to short internal branches difficult to resolve, and therefore we consider that additional phylogenetically informative molecular markers and a thorough morphological examination are necessary requirements to take any taxonomic action.

Our study further highlights the importance of evaluating different phylogenetic reconstruction methods to avoid systematic classifications that reflect incongruent or inconsistent species trees and that might lead to conflictive diagnostic characters, confounding taxonomic efforts.

## Acknowledgements

We are extremely thankful to James Morin, Gene Helfman, Masanori Sato and Marcelo Fukuda for providing specimens collected in different locations around the world. We want to thank Juliette Gorson for her help with phylogenetic analyses and two talented high school students, Jacob Djibankov and Debjani Das, for assisting with lab work. This research was funded through the Systematics Research Fund of The Linnaean Society of London, a Mini-ARTS Award from the Society of Systematic Biologists and a Science Scholarship from The Graduate Center of the City University of New York to AV. MH acknowledges funding from the Camille and Henry Dreyfus Teacher-Scholar Award and NSF awards CHE-1247550 and CHE-1228921.

